# OmpK36 and TraN facilitate conjugal transfer of the *Klebsiella pneumoniae* carbapenem resistance plasmid pKpQIL

**DOI:** 10.1101/2020.07.01.180638

**Authors:** Wen Wen Low, Joshua LC Wong, Alejandro Peña, Chloe M Seddon, Tiago RD Costa, Konstantinos Beis, Gad Frankel

## Abstract

We investigated the mechanism of conjugal transfer of the endemic *Klebsiella pneumoniae* carbapenem resistance plasmid, pKpQIL. Transfer efficiency of this plasmid was found to be dependent on the expression of the major outer membrane porin, OmpK36, in recipient cells. We also found that conjugal uptake is reduced in recipients expressing an OmpK36 isoform associated with the globally pervasive *K. pneumoniae* ST258 clade (OmpK36^ST258^). This reduction was attributed to a glycine-aspartate insertion in loop 3 of OmpK36^ST258^, which constricts the pore by 26%. Deletion of *finO*, which encodes an RNA-binding protein, derepressed transfer of pKpQIL and enabled visualisation of the conjugation pilus and OmpK36-dependent conjugation in real time. While deletion of *traN* abolished pKpQIL conjugation, substituting *traN* in pKpQIL with its homologue from R100-1 circumvented OmpK36 dependency. These results suggest that OmpK36 in recipient *K. pneumoniae* and the pKpQIL-encoded TraN in donor bacteria cooperate to facilitate plasmid transfer. This is the first report since 1998 to suggest a novel recipient cell receptor for IncF plasmid transfer and supports the idea that TraN mediates receptor specificity for plasmids belonging to this incompatibility group.

## Introduction

Conjugation, discovered by Lederberg and Tatum in 1946, refers to the contact-dependent unidirectional transfer of genetic material between donor and recipient bacteria ^1^. While the structures of the F-plasmid-encoded type IV secretion system (T4SS) and pilus were recently elucidated by cryoelectron tomography and cryoelectron microscopy reconstructions, respectively ^2,3^, the mechanism of conjugal DNA transfer remains largely unknown. However, with the increasing threat to public health from the spread of antimicrobial resistance, it is fundamentally important that we close this gap in knowledge as conjugation is at the heart of the spread of resistance genes amongst key human pathogens ^4^.

*Klebsiella pneumoniae* is a successful modern pathogen which thrives in the nosocomial environment ^5^. Carbapenem resistant *K. pneumoniae* (CRKP) have been associated with a multitude of conjugative plasmids encoding carbapenemases. In particular, the globally pervasive sequence type ST258 is largely associated with the *bla*^KPC^-encoding IncFII^K^ pKpQIL-family plasmids ^6^. These plasmids have been linked to outbreaks of CRKP in Europe, Asia and the United States ^7,8^. Other carbapenemases include *bla*^OXA-48-like,^ which is commonly associated with the IncL/M plasmid pOXA-48a, and *bla*^VIM^ and *bla*^NDM^, which are found on a diverse range of plasmids ^9^.

Plasmid-encoded carbapenem resistance in *K. pneumoniae* is often augmented by chromosomally-encoded mutations in the outer membrane (OM) porins OmpK35 and OmpK36 ^10^. Many clinical ST258 isolates express a variant of OmpK36 containing a glycine-aspartic acid (GD) insertion in loop 3 (L3) ^11^. We have recently shown that this insertion results in a 26% reduction in the pore diameter and a 4-fold increase in the minimum inhibitory concentration of meropenem ^12^.

The transfer of single stranded DNA by conjugation involves three main components: a T4SS, a coupling protein (T4CP) and a relaxosome ^13,14^. T4SSs are highly diverse, and can be classified into F-type, P-type and I-type. Studies on the mechanism and structural basis of conjugation have largely been performed in *Escherichia coli* K12 using the F-plasmid (F-type) or the IncW plasmid, R388 (P-type) ^2,15^. Consequently, little is currently known about conjugal transfer of contemporary plasmids that circulate in hospital environments, e.g. pKpQIL. In this study, we investigated the relationship between two key determinants of carbapenem resistance, by determining the role of OmpK36 in the conjugal transfer of pKpQIL.

## Results

### Conjugal uptake of pKpQIL is dependent on OmpK36 in recipient *K. pneumoniae*

We first determined if OmpK36 is essential for pKpQIL conjugation using *K. pneumoniae*Δ*ompK36* as a recipient. This revealed that conjugation frequency of pKpQIL was approximately 100-fold higher into wild type (WT) *K. pneumoniae* recipients, expressing OmpK36, compared to *K. pneumoniae*Δ*ompK36* (Fig. 1A). We next investigated if conjugal transfer of pKpQIL may also be affected by the relative abundance of OmpK36 within the OM. OmpK36 is homologous to the *E. coli* OM porin OmpC, which is osmoregulated through the EnvZ/OmpR two-component regulatory system ^16^. Thus, we modulated OmpK36 expression by growing *K. pneumoniae* recipients in media containing different concentrations of sodium chloride (NaCl). Recipients were then mixed with pKpQIL-carrying donor bacteria and the mixtures were allowed to conjugate on agar containing the same NaCl concentration that the recipient cells had been cultured in. We found a significant reduction in conjugation frequency of pKpQIL as NaCl concentration increased from 0.5 g/L to 20 g/L (Fig. 1B). Using a transcriptional reporter (Fig. S1) in which superfolder GFP (sfGFP) was expressed from the *ompK36* promoter revealed that, contrary to *ompC* in *E. coli* ^16^, there was reduced promoter activity in response to increasing medium osmolality (Fig. 1C); we observed no change in expression using the constitutive Biofab promoter-driven sfGFP as a control (Fig. S1). Moreover, western blotting of N-terminal His-tagged OmpK36 (inserted downstream of the signal peptide) revealed reduced abundance of OmpK36 in response to increasing medium osmolality (Fig. 1D & E). Therefore, we concluded that conjugal uptake of pKpQIL is dependent on both the presence and the relative abundance of OmpK36 within the bacterial OM of the recipient.

**Figure 1.**
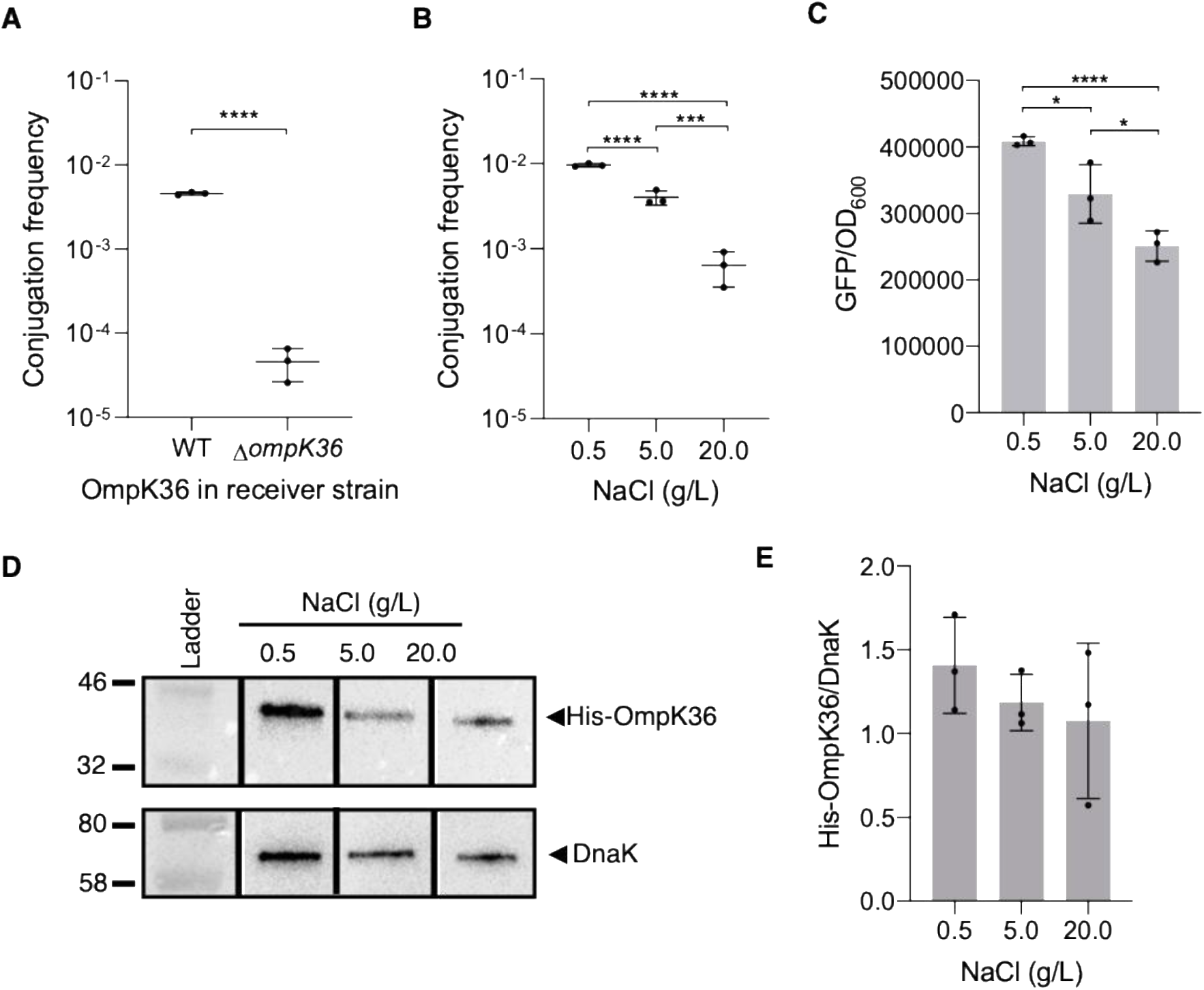
Conjugal transfer of pKpQIL is dependent on OmpK36. Conjugation frequency of pKpQIL was compared (**A**) between recipients expressing or missing OmpK36, and (**B**) between OmpK36-expressing recipients grown at different NaCl concentrations. Conjugation frequency was determined as CFU transconjugants/ CFU recipients. (**C**) Activity of the OmpK36 promoter, p*ompK36*, was measured by GFP emission from strains containing a synthetic p*ompK36*-sfGFP reporter. GFP emission was normalized to OD^600^. * P ≤0.05, *** P ≤ 0.001, **** P ≤ 0.0001. (**D** and **E**) Expression of OmpK36 at different NaCl concentrations was determined by Western blot (anti-His) and densitometric analysis with normalization to a loading control, DnaK. A representative blot is shown in (**D**) and densitometric quantification from three biological replicates is shown in (**E**).

### The L3 GD insertion in OmpK36^ST258^ inhibits conjugal transfer of pKpQIL

As an OmpK36 variant has been associated with the highly resistant ST258 clade carrying pKpQIL, we tested the hypothesis that conjugation frequency of pKpQIL may be higher in recipients expressing this resistant isoform. This variant, OmpK36^ST258^, when compared to an isoform from the prototype *K. pneumoniae* strain ATCC43816, herein referred to as OmpK36^WT^, contains the L3 GD insertion and a loop 4 (L4) leucine-serine-proline (LSP) insertion (Fig. 2A) ^12^.

**Figure 2.**
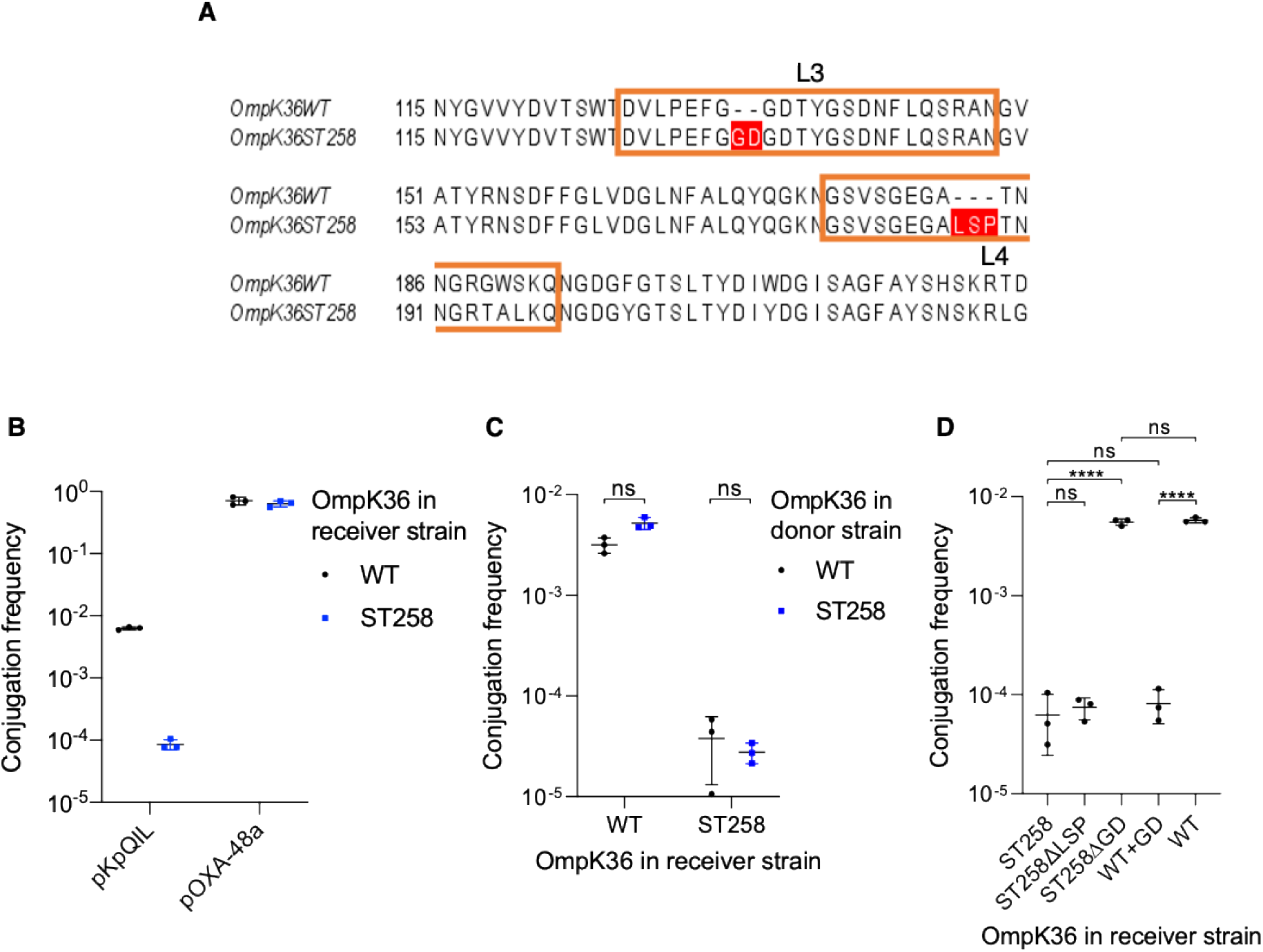
The L3 GD insertion in OmpK36^ST258^ results in 100-fold lower conjugal uptake of pKpQIL in recipient cells. (**A**) Amino acid sequence alignment of OmpK36^WT^ (Genbank accession no: AIK83412.1) and OmpK36^ST258^ (Genbank accession no: WP_00415112) shows the glycine-aspartic acid (GD) insertion in L3, and the leucine-serine-proline (LSP) insertion in L4 of OmpK36^ST258^. (**B**) Conjugation frequency of pKpQIL and pOXA-48a into recipients expressing either the WT or ST258 isoform of OmpK36. (**C**) Conjugation frequency of pKpQIL from donor cells expressing either OmpK36^WT^ or OmpK36^ST258^. (**D**) Conjugation frequency of pKpQIL into recipients expressing different OmpK36 isoforms. Conjugation frequency was determined as CFU transconjugants/ CFU recipients. **** P ≤ 0.0001, ns = non-significant.

Comparing the conjugal uptake of pKpQIL between recipient cells expressing either OmpK36^WT^ or OmpK36^ST258^ unexpectedly revealed that recipients expressing OmpK36^ST258^, similar to the *K. pneumoniae*Δ*ompK36* recipients (Fig. 1A), displayed 100-fold lower conjugal uptake of pKpQIL compared to recipients expressing OmpK36^WT^ (Fig. 2B). Previous work has shown that growth of these isogenic recipients shows no significant difference in rich media ^12^. We also determined if expression of OmpK36^ST258^ impairs conjugal uptake in other carbapenemase-encoding plasmids and assessed this using pOXA-48a, an IncL/M carbapenem resistance plasmid, which is transferred via an I-type T4SS ^17^. In contrast to pKpQIL, there was no significant difference in the conjugation frequency of this plasmid into recipients expressing either OmpK36^ST258^ or OmpK36^WT^ (Fig. 2B). Furthermore, there is no significant difference in pKpQIL conjugation frequency when OmpK36^ST258^ was expressed in donor cells (Fig. 2C). Thus, transfer of pKpQIL is specifically affected by expression of the OmpK36^ST258^ variant in recipient bacteria.

To determine if the ST258-associated insertions in OmpK36 contribute to the inhibition of conjugal uptake of pKpQIL, we measured conjugation frequency into strains expressing mutant isoforms of OmpK36^ST258^ (Fig. 2D). We first assessed the effect of the LSP insertion found in the surface exposed L4 of OmpK36^ST258^. This revealed no significant difference in conjugation frequency between recipients expressing OmpK36^ST258^ or OmpK36^ST258ΔLSP^ (Fig. 2D). In contrast, assessing the effect the L3 GD insertion has on conjugation frequency revealed that recipients expressing OmpK36^ST258ΔGD^ were 100-fold more permissive for conjugation compared to recipients expressing OmpK36^ST258^. This effect was reconstituted by inserting GD into L3 of OmpK36^WT^ to generate OmpK36^WT+GD^, as there was no significant difference in conjugation frequency between recipients expressing OmpK36^WT+GD^ and OmpK36^ST258^. This suggests that the L3 GD insertion found in OmpK36^ST258^ affects conjugal uptake of pKpQIL.

We have recently shown that that the GD insertion results in a conformational change to L3 that is stabilised by a salt bridge formed between D114 and R127 at the barrel face of the pore ^12^. To determine if this salt bridge affects plasmid transfer, we compared conjugation frequency into recipients expressing OmpK36^ST258^ and OmpK36^ST258R127A^ but found no significant difference (Fig. S2). We further hypothesised that inhibition of plasmid transfer may be due to the introduction of a negative charge associated with the GD insertion. To test this, we generated a D116N mutant (OmpK36^ST258D116N^). However, we observed no significant difference in conjugation frequency between recipients expressing OmpK36^ST258D116N^ or OmpK36^ST258^ (Fig. S2). These experiments suggest that neither the salt bridge nor the charge associated with the GD insertion affect conjugal uptake of pKpQIL.

### L3 GD insertion conjugal transfer inhibition in real time

Neither OmpK36 in *K. pneumoniae*, nor OmpC in *E. coli*, have previously been implicated in conjugation. Furthermore, as the L3 GD insertion, which interferes with conjugation, is concealed inside the pore and is not surface exposed, it was necessary to validate our conclusions using an alternative methodology. To this end, we developed a real time, fluorescence-based, conjugation assay, for which pKpQIL was tagged with sfGFP under the control of a *lac* promoter at the disrupted *aadA* gene, to generate pICC4000. pICC4000 was then transferred into a recombinant *K. pneumoniae* donor, we named GFP donor (GFP-D), carrying chromosomal *lacI* expressed under the control of the constitutive promoter Biofab, which represses sfGFP expression. When pICC4000 is conjugated into a recipient *K. pneumoniae* lacking recombinant *lacI*, sfGFP is expressed and the resulting fluorescence is used as an indirect measure of conjugation (Fig. 3A). However, we were unable to detect an increase in fluorescence over 6 h of conjugation even with *K. pneumoniae* recipients expressing OmpK36^WT^, which suggested that the assay might not be sensitive enough to detect pKpQIL transfer.

**Figure 3.**
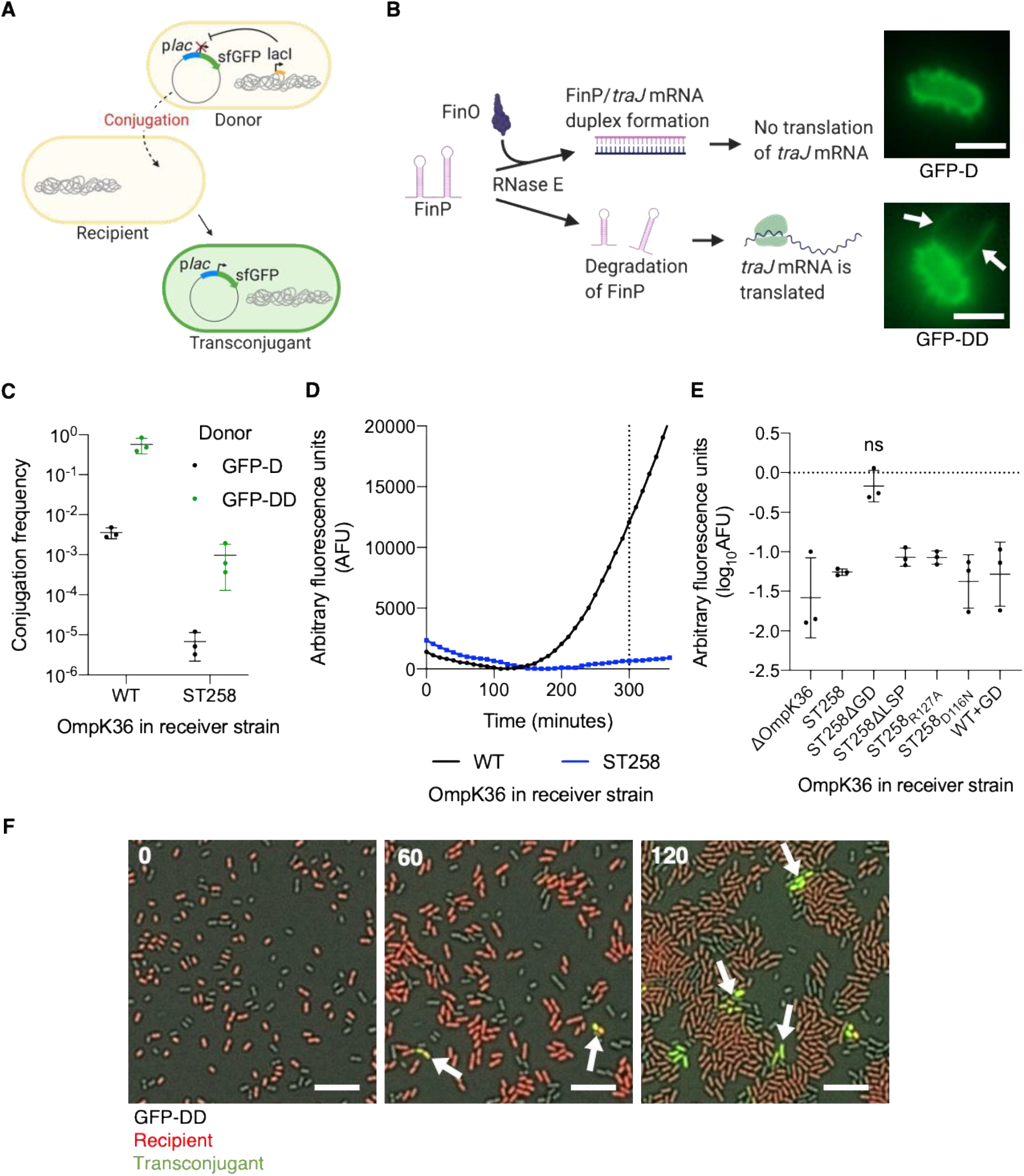
Real time visualisation of conjugation inhibition by the L3 GD insertion. (**A**) Schematic representation of the synthetic conjugation system of the sfGFP-tagged pKpQIL. (**B**) Schematic representation of the regulatory role of FinO in pKpQIL transfer. FinO protects FinP antisense RNA from degradation by RNase E, leading to the formation of a FinP/*traJ* mRNA duplex that prevents translation of *traJ* mRNA. Conjugal transfer can be derepressed by deleting *finO* resulting in degradation of FinP and the translation of *traJ* mRNA. Immunofluorescence images of the repressed donor, GFP-D, and the derepressed donor, GFP-DD, using anti-pilus antibodies indicate the presence of conjugative pili (indicated with arrows) on GFP-DD but not GFP-D cells. Scale bar = 2 μM. (**C**) Plasmid transfer from GFP-D and GFP-DD into recipients expressing either OmpK36^WT^ or OmpK36^ST258^. Conjugation frequency was determined as CFU transconjugants/CFU recipients. (**D**) Plasmid transfer from GFP-DD was monitored in real time by measuring GFP emission over 6 h into recipients expressing either OmpK36^WT^ or OmpK36^ST258^. Arbitrary fluorescence units (AFU) were normalised to the minimum GFP emission recorded for each conjugation mixture. (**E**) The log-fold difference in AFU recorded at *t* = 300 min respective to the OmpK36^WT^-expressing recipient strain was calculated for recipients expressing various isoforms of OmpK36. ns = non-significant. (**F**) Still images from live microscopy of conjugation mixtures containing GFP-DD donor cells, *tdTomato*-expressing recipient cells (red), and transconjugant cells (green) at 0, 60 and 120 min. New clusters of transconjugant cells are indicated with arrows. Scale bar = 10 μM. The complete time series can be viewed as a movie (**Supplementary movie**).

Due to a disruption in the conjugation inhibitor gene, *tir* ^18^, pOXA-48a conjugates at frequencies 100-fold higher than pKpQIL (Fig. 2C). We therefore used a similarly tagged pOXA-48a (pICC4001), to evaluate the functionality of this assay in a derepressed system. Using pICC4001 in this setup, we observed an increase in arbitrary fluorescence units (AFU) over 6 h of conjugation into *K. pneumoniae* recipients expressing either OmpK36^ST258^ or OmpK36^WT^ (Fig. S3A & B). This suggests that the low frequency of pKpQIL transfer hindered detection of sfGFP in our setup. To overcome this inherent limitation in detecting pKpQIL conjugal transfer, we deleted the negative regulator of conjugation, *finO*, from pKpQIL to generate pKpQILΔ*finO* (pICC4002) (Table S1). FinO represses expression of the *tra* operon by binding and stabilising FinP, which inhibits expression of the transcription factor TraJ ^19^ (Fig. 3B). Growth of *K. pneumoniae* carrying pICC4002 was attenuated in both rich and minimal media compared to a strain carrying WT pKpQIL (Fig. S4A & B), suggesting that derepression comes at a fitness cost. To confirm derepression of the *tra* operon, we visualised *K. pneumoniae* donor strains carrying either pKpQIL or pICC4002 by immunofluorescence microscopy using rat polyclonal antibodies made against the purified pKpQIL-encoded pilus (Fig. S4C). This revealed that *K. pneumoniae* carrying pICC4002 expressed conjugative pili constitutively, while no pili were seen in *K. pneumoniae* carrying pKpQIL (Fig. 3B). Consistently, there is a 100-fold increase of conjugation frequency of pICC4002 into recipient cells. Importantly, the difference in conjugation frequency into recipients expressing the different OmpK36 isoforms seen for pKpQIL was maintained for pICC4002 (Fig. 3C).

To assess inhibition of transfer by the L3 GD insertion in real time, pKpQILΔ*finO* was tagged with p*lac*-sfGFP (pICC4003) (Table S1) and mobilized into the *lacI*-expressing strain to generate the GFP derepressed donor (GFP-DD). Using this donor, we saw an increase in AFU over 6 h for recipients expressing OmpK36^WT^ but not OmpK36^ST258^ (Fig. 3D). Repeating the assay with the various OmpK36 isoforms revealed that only recipients expressing OmpK36^ST258ΔGD^ showed no significant log-fold difference in AFU compared to recipients expressing OmpK36^WT^, which validates the conclusion that the L3 GD insertion leads to inhibition of conjugal uptake of pKpQIL (Fig. 3E). The GFP-DD donor strain also allowed us to observe pKpQIL transfer into recipient cells through live microscopy. In this setup, recipients constitutively expressing tdTomato begin to co-express sfGFP upon acquisition of pICC4003 (Fig. 3F and Video). Taken together, these results suggest that OmpK36 is a conjugation receptor of pKpQIL in recipient *K. pneumoniae* and that the L3 GD insertion interferes with this activity.

### TraN cooperates with OmpK36 during conjugation of pKpQIL

In F-plasmid, TraN was suggested to play a role in mating pair stabilisation (MPS) via interactions with OmpA in *E. coli* K12 recipient bacteria ^20^. While direct TraN:OmpA interactions have not been demonstrated, their cooperation has been shown to be plasmid-specific as substituting the F-plasmid *traN* with *traN* from another IncF plasmid, R100-1, abrogated OmpA-dependency ^21^. Based on this, we hypothesised that TraN could be the pKpQIL-encoded factor in donor cells mediating the OmpK36-dependency of transfer into *K. pneumoniae* recipient bacteria.

To test this hypothesis, we first deleted *traN* from pKpQIL to generate pKpQILΔ*traN*, (pICC4004) (Table S1); conjugative transfer of pICC4004 into WT *K. pneumoniae* recipient cells was reduced to below the limit of detection (Fig. 4A), implying that TraN plays an essential role in pKpQIL transfer. Next, as TraN of R-100 bypassed OmpA dependency in F-plasmid conjugation, we substituted *traN* of pKpQIL with *traN* of R100-1, to generate pKpQIL^TraN_R100-1^ (pICC4005) (Table S1). Sequence alignment of TraN from pKpQIL and R100-1 revealed that the C-terminal region is largely conserved. The central region of TraN from pKpQIL located between aa 171 and aa 337 is more divergent and includes several large insertions (Fig. S5). This more divergent central region is similar to what was observed in sequence alignments between TraN from F-plasmid and R100-1 ^22^. Despite these differences in amino acid sequence, conjugation using a donor strain carrying pICC4005 revealed that TraN from R100-1 was able to complement the loss of expression of TraN from pKpQIL (Fig. 4B). Importantly, there was also no significant difference in conjugation frequency between recipients expressing different OmpK36 isoforms and plasmid transfer was no longer affected by the L3 GD insertion (Fig. 4B). This was further validated using the real time assay where the AFU over time was seen to increase in all recipient strains tested and there was no significant difference in log^10^AFU measured at 300 min for all recipients (Fig. 4C and 4D). This suggests that TraN mediates OmpK36-dependency in pKpQIL conjugation in *K. pneumoniae*.

**Figure 4.**
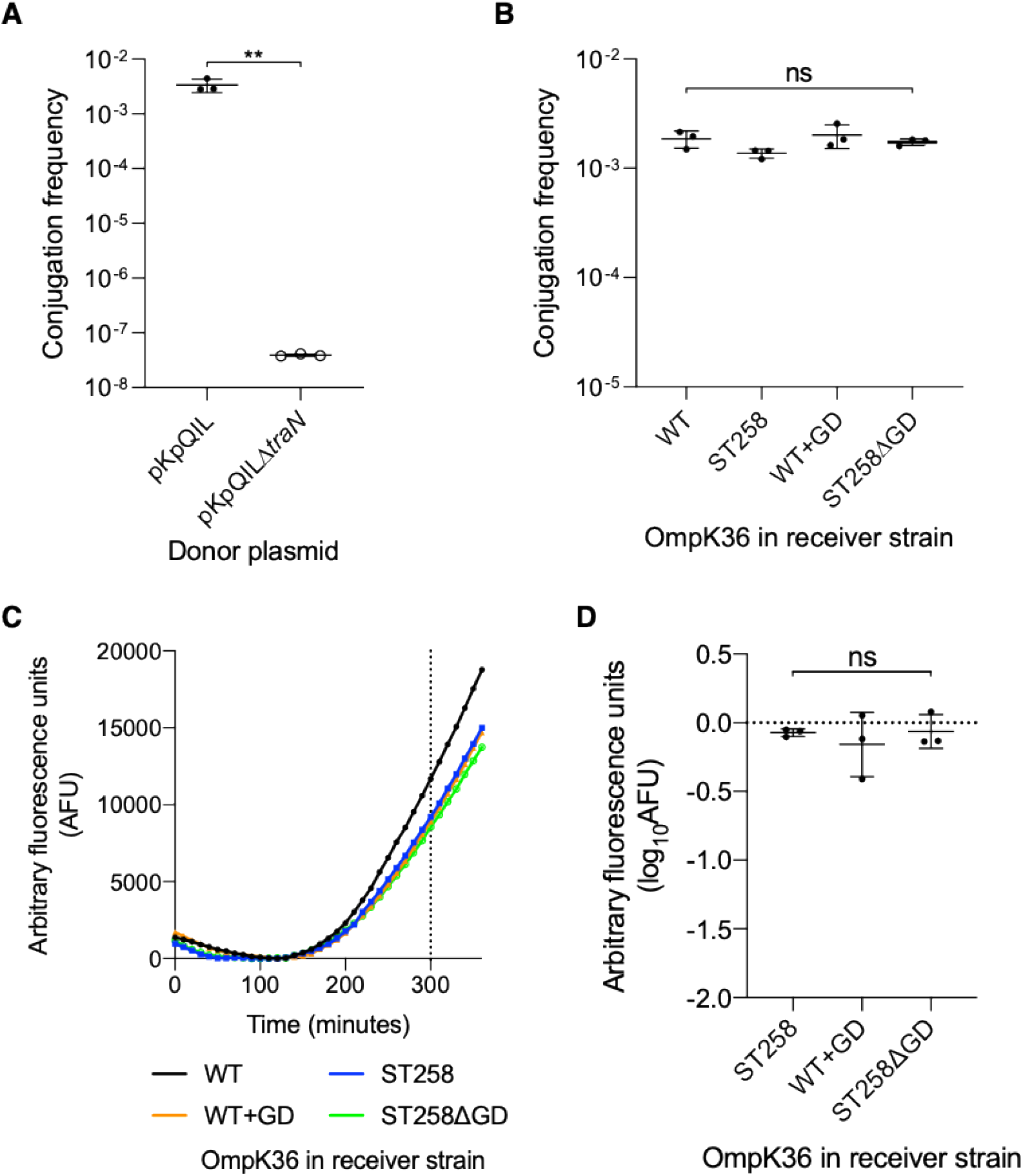
TraN in pKpQIL mediates OmpK36-dependency during conjugative transfer. (**A**) Conjugation frequency of pKpQIL and pKpQILΔ*traN* into OmpK36^WT^-expressing recipient cells. Open circles indicate that no transconjugants were observed at the limit of detection. (**B**) Conjugation frequency from a donor strain carrying pKpQIL^TraN_R100-1^ into recipients expressing various OmpK36 isoforms. Conjugation frequency was determined as CFU transconjugants/CFU recipients. (**C**) Transfer of pKpQIL^TraN_R100-1^ was monitored in real time using the fluorescence-based assay. Arbitrary fluorescence units (AFU) presented have been normalized to the minimum GFP emission recorded for each conjugation mixture. (**D**) The log-fold difference in AFU recorded at *t* = 300 min respective to the OmpK36^WT^-expressing recipient strain was calculated for each recipient. ** P ≤0.01, ns = non-significant.

## Discussion

In this study we show that OmpK36 in *K. pneumoniae* recipient cells mediates conjugal uptake of pKpQIL. Mechanistically, both reduced expression of OmpK36 and a constriction of its pore by a GD insertion in L3 reduces conjugal uptake into recipient cells. We excluded the possibility that the charge introduced by the GD insertion, which is concealed inside the pore, or salt bridge formed between D114 and R127 contributes to conjugal inhibition. As disrupting the salt bridge does not alter the permeability of OmpK36^ST258^, our data suggests that pore diameter is a key parameter affecting conjugation efficiency ^12^.

Importantly, OmpK36-dependency appears to be specific to pKpQIL and substituting *traN* in pKpQIL with its homologue from R100-1 abrogates this dependency. This is reminiscent of OmpA-dependency observed with F-plasmid transfer in *E. coli* K12 where R100-1 transfer was also unaffected by mutations in OmpA ^22^. Overall, this provides compelling evidence that conjugation receptors are highly specific to the plasmids being transferred even if they are from the same incompatibility groups and express machinery of the same type classification. We assume that receptor proteins are even more diverse for plasmids transferred using other T4SS types. This may explain why studies which have tried to determine conjugation receptors for other conjugative plasmids, like the IncW plasmid R388 which transfers via a P-type T4SS, failed to identify OmpA in their screens ^23^.

TraN is an IncF plasmid-specific conjugation component which plays a role in MPS. This process involves the formation of multicellular mating aggregates that increases plasmid transfer efficiency. In F-plasmid, the functionality of TraN was studied using a chemically-induced amber stop codon mutant. Plasmids expressing this TraN mutant retain F pilus expression and were able to conjugate on solid media ^20^. Here, we demonstrated that TraN mediates OmpK36-dependency in pKpQIL transfer and that a *traN* deletion abrogated conjugation on solid media. Biochemical analysis of conjugation transfer subunits from F-plasmid found that TraN is localised to the OM but might not be part of the core conjugation machinery ^2^. Therefore, our data supports a model whereby cooperation between OM proteins in the donor and recipient cells are essential for DNA transfer. Importantly, deletion of *traN* has a much greater effect on conjugation than deletion of *ompK36*. This suggests that OmpK36 may not be the only recipient cell outer membrane protein involved in mediating pKpQIL conjugation. On the other hand, TraN, as expressed in this system appears to be an essential component that not only plays a role in MPS but potentially serves as a structural T4SS protein. Therefore, an important question for future studies is the functional and structural relationships between the T4SS, the MPS complex (e.g. TraN:OmpK36) and the conjugation pilus.

In addition, we developed an assay that allowed us to study conjugative transfer of pKpQIL in real time and in the absence of selection pressure. As pKpQIL conjugates naturally at very low frequencies (<1%), we deleted the conjugation inhibitor gene, *finO*. Using immunofluorescence microscopy with antibodies raised against the conjugative pili, we did not observe piliation on the surface of donor cells carrying pKpQIL. Instead, piliation was observed on donor cells carrying pKpQILΔ*finO*. This suggests that the expression of pKpQIL transfer genes are tightly regulated and that assembly of the conjugation machinery may only be triggered in the presence of a signal that is yet to be determined. Deleting *finO* disrupts regulation resulting in constitutive expression and assembly of the conjugation machinery even in the absence of recipient cells.

De-repression of pKpQIL results in an overall 100-fold increase in conjugation frequency into recipient cells regardless of the OmpK36 isoform being expressed. We predict that it is unlikely that downstream processes like DNA processing and plasmid recircularisation are affected in the OmpK36^ST258^-expressing recipients. Therefore, the processes that are most likely to be affected by the GD insertion are the initial formation of contacts between donor and recipient cells and the DNA transfer process itself. As the L3 GD insertion is not surface exposed, OmpK36 may have a dual role as a receptor for conjugation via an interaction with TraN, and as a conduit through which plasmid ssDNA passes to traverse the recipient cell OM. Although not conforming to the current models, the role of OmpK36 as a pore through which DNA traverses the OM cannot be completely discounted, especially as the mutation affecting conjugation efficiency greatly constricts the porin channel.

## Acknowledgments

We would like to thank Fernando de la Cruz for sharing the R100-1 and pSU2007::Tnlux plasmids. We would also like to acknowledge Claire Elek and Michelle Yeap for their assistance in generating strains used in this study. This project has been supported by grants from the Wellcome Trust, the MRC and the BBSRC.

## Author Contributions

W.W.L., J.L.C.W., A.P. and C.M.S conducted the experiments.

W.W.L., J.L.C.W., and G.F. designed the experiments

W.W.L., J.L.C.W., T.R.D.C., K.B., and G.F. participated in supervision

W.W.L., and G.F. wrote the paper

## Methods

### Bacterial strains and plasmids

The bacterial strains (S), vectors (V) and primers (P) used in this work are listed in Supplementary Tables 2, 3 and 4 respectively. Unless otherwise stated, bacteria were cultured in Luria Bertani (LB) broth at 37°C, 200 RPM. When needed, antibiotics were used at the following concentrations: ertapenem (0.5 μg/ml), streptomycin (50 μg/ml), kanamycin (50 μg/ml), chloramphenicol (30 μg/ml), gentamicin (10 μg/ml).

Unless otherwise stated, ICC8001, a rifampicin resistant derivative of *K. pneumoniae* ATCC43816 ^12^, served as the donor for all conjugative plasmids used in this study. To generate the different ICC8001 donor strains, plasmids were conjugated from strains S3 and S4 (Table S2) into ICC8001 containing pACBSR (SmR) and selected on the appropriate antibiotics. DNA transfer was confirmed by PCR and pACBSR was cured by serial passaging.

Genomic insertions, *ompK36* substitutions, and plasmid variants were generated using a two-step recombination methodology as previously described ^24^. All genomic insertions were targeted to the 3’ end of the *glmS* gene of ICC8001 and all constructs were generated by Gibson Assembly (New England Biolabs, US).

For construction of the *ompK36* transcriptional reporter, a 400 bp fragment preceding the start codon of *ompK36* was amplified from ICC8001 with P1/2. The fragment includes all known binding sites of the osmoregulatory transcription factor, OmpR, as identified in Tsai et al, 2011 ^10^. This was assembled into linearized V3 (P3/4) to generate V4. The N-terminal 6xHis-tagged OmpK36 was generated by site directed mutagenesis of V5 with P5/6 to generate V6.

All isogenic OmpK36 recipient strains are a derivative of ICC8003 (S2), which itself was generated from ICC8001 ^12^. OmpK36^ST258D116N^ was generated by site-directed mutagenesis of V7 with P7/8 and inserted into a Δ*ompK36* mutant of ICC8003.

To generate a constitutive *lacI*-expressing donor strain, *lacI* was amplified using pET-28a as a template with P9/10 and assembled into linearized V3 (P11/12) to generate V9. To generate *tdTomato*-expressing recipients, *tdTomato* was amplified sing pCSCMV-tdTomato as a template with P13/14 and assembled into linearized V3 (P11/12) to generate V11.

To generate the p*lac*-sfGFP tagged pKpQIL (pICC4000), sfGFP was amplified from V3 (P15/16) and the *lac* promoter was amplified from V12 (P17/18). 500 bp upstream and downstream flanking the insertion site in the disrupted *aadA* gene was amplified from pKpQIL (P19/20 and P21/22). All fragments were assembled into linearized V1 (P23/24) to generate V13. To generate the tagged pOXA-48a, the p*lac*-sfGFP fragment (P25/26), and 500 bp upstream and downstream flanking the insertion site in the disrupted *tir* gene in pOXA-48a (P27/28 and P29/30) were amplified and assembled into linearized V1 to generate V14.

The *finO* deletion vector (V15) was generated by assembling 500 bp upstream and downstream flanking regions of the gene (P31/32 and P33/34) into linearized V1. The *traN* deletion vector (V16) was generated by assembling 500 bp upstream and downstream flanking regions of *traN* from pKpQIL (P35/36 and P37/38) into linearized V1. The *traN* substitution vector (V17) was generated by amplifying *traN* from R100-1 (P39/P40) and assembling it into linearized V16 (P41/42).

### Selection-based conjugation assays

All recipient strains used in these assays were transformed with pSEVA471, a low-copy number plasmid encoding streptomycin resistance. For all experiments, overnight cultures of donor and recipient bacteria were washed in phosphate-buffered saline (PBS). For pKpQIL and pOXA-48a conjugation mixtures, donor and recipient cells were mixed at a ratio of 8:1 and diluted in PBS (1 in 25 v/v). For surface mating experiments, 40 μl of the final conjugation mixture was spotted onto LB agar plates and incubated for 6 h at 37°C. The spots were collected and resuspended in 1 ml of sterile PBS for serial dilution. Recipient colonies were selected on streptomycin-containing LB agar plates. Transconjugants were selected on plates supplemented with streptomycin and ertapenem. Conjugation frequency was given as the CFU of transconjugants/CFU of recipients.

### Real-time conjugation assays

Conjugation mixtures were prepared by mixing washed overnight cultures of donor and recipient bacteria. It was determined that maximal fluorescence emission was obtained when donor and recipient bacteria were mixed at a 1:1 ratio without dilution (pKpQIL) and with 1 in 25 v/v dilution in PBS (pOXA-48a). 8 μl of the conjugation mixture was spotted onto 300 μl of LB agar in a 96-well black microtiter plate in technical triplicate. The plates were incubated for 6 h at 37°C with fluorescence readings taken at 10-min intervals on a FLUOstar Omega (BMG Labtech, UK). Arbitraty fluorescence units (AFU) are normalized to the minimum GFP emission recorded for each sample. The fold difference in AFU at *t* = 300 min for each mutant recipient strain (*x*) was calculated as log^10^ (AFU^X^ / AFU^WT^).

### Agarose pad live microscopy

Bacterial conjugation was visualized over time on a Celldiscoverer 7 live cell imaging microscope (Zeiss, Germany). For these experiments, the GFP-DD donor strain and *tdTomato-*expressing (ICC8003) recipients were used. Overnight cultures of donor and recipient bacteria were washed in PBS and mixed in a 1:1 ratio and 8 μl spotted onto a 1 cm^2^ 2% agarose (w/v) pad supplemented with M9 salts and 0.4% glucose (w/v). The pad was inverted into a μ-Slide 2-well fitted with a polymer coverslip (Ibidi, Germany). The sample was maintained at 37°C throughout live imaging. Images were acquired every 10 min for 3.5 h and processed using Zen 2.3 (Blue Version; Zeiss, Germany).

### Purification of conjugative pili and generation of polyclonal antibodies

2 L of ICC8001pKpQILΔ*finO* overnight cultures were harvested by centrifugation at 7000 x *g* for 20 min and resuspended in 40 ml of cold 1X PBS. The resuspended cells were passed through a 25G needle 30 times. “Shaved” bacteria were centrifuged at 50000 x *g* for 1 h. The supernatant was mixed with 5% PEG 6000 with constant stirring for 1 h at 4°C. Conjugative pili were precipitated by centrifugation at 50000 x *g* for 30 min. The pellet was resuspended in a buffer containing 50 mM Tris pH8, 1 M NaCl and dialysed overnight against the same buffer. The purified pili were visualised by negative stain electron microscopy to assess for pilus integrity and purity. Rat polyclonal antibodies were raised against the purified pili (ThermoFisher Scientific, US).

### Growth curves

Saturated overnight cultures were diluted 1:100 in 200 μl of fresh rich LB or minimal M9 media supplemented with glucose at 0.4% w/v. OD600nm readings were taken on a FLUOStar Omega (BMG Labtech, UK) at 30 min intervals with shaking at 200 rpm at 37 °C between readings.

### Immunofluorescence microscopy

Strains used for immunofluorescence microscopy were grown overnight in LB without shaking at 37°C. Overnight cultures were diluted 1 in 20 (v/v) in fresh LB and 80 μl was spotted onto glass coverslips before incubation at 37°C for 1.5 h. Excess media was removed, and coverslips were allowed to dry fully before fixation in 4% paraformaldehyde for 20 min at room temperature. Fixed samples were washed in PBS and blocked in 2% bovine serum albumin (BSA) in PBS (w/v). Samples were washed three times before incubation with anti-pili rat polyclonal antibodies (1:500 dilution) for 1 h at room temperature. Samples were washed three times in PBS and incubated with Alexa Fluor 488 Donkey anti-rat IgG antibodies (Jackson Immunoresearch, 1:200 dilution) for 1 h at room temperature. Coverslips were washed three times in PBS and mounted onto slides using VECTASHIELD® Hardset™ Antifade Mounting Medium with DAPI according to the manufacturer’s instruction. Slides were analysed using a 100X objective lens on a Zeiss Axio Observer 7 microscope and images were processed on Zen 2.3 (Blue Version; Zeiss, Germany).

### OmpK36 promoter activity assay

For measuring promoter activity, the promoter of *ompK36* or the constitutive Biofab promoter were fused to a gene encoding sfGFP and inserted into the genome of ICC8001. Medium osmolality was modulated by changing the concentration of sodium chloride (NaCl) in low salt LB. Overnight cultures of the p*ompK36* and Biofab reporter strains were grown in different osmolality media and normalized to an OD^600^ of 0.5 before fluorescence was measured.

### Western immunoblotting

ICC8001 expressing an N-terminal 6XHis-tagged OmpK36 was cultured overnight in LB media of different osmolalities. 1 ml of overnight culture adjusted to an OD^600^ of 0.5 was pelleted and cells were lysed by sonication. Total protein amount in sonicated whole cell lysates were quantified on a Qubit™ 4 Fluorometer (ThermoFisher Scientific, US). 5 μg of cell lysate was resuspended in 2X Laemlli buffer (from 5X stock, 312.5 mM Tris-HCl, pH 6.8, 10% w/v SDS, 50% v/v glycerol, 0.5 M DTT, 0.05% w/v bromophenol blue). Samples were boiled at 100°C for 5 min prior to SDS-PAGE separation on 12% acrylamide Mini-protean TGX precast gels (Bio-Rad Laboratories, US) and Western blot analysis.

### Statistical analysis

All statistical analysis was performed on Prism 8 (GraphPad Software Inc, US). All data is presented as the mean of three biological replicates ± standard deviation. Conjugation frequencies were compared between recipients by an unpaired *t*-test or, where more than 2 recipients were being compared, by ordinary one-way analysis of variance (ANOVA) correcting for FDR using the Benjamini, Krieger and Yekutieli method. P values <.05 were considered significant and the desired false discovery rate was set to 0.05.

### Multimedia File

#### Live microscopy of pKpQIL conjugation

The conjugation mixture contains GFP-DD donor cells, *tdTomato*-expressing recipient cells (red), and transconjugants cells (green). Each frame was captured 10 min apart. The video plays at 2400X actual speed.

## References

1. Lederburg, J. & Tatum, E. L. Gene Recombination in Escherichia coli. Nature 158, 558 (1946).

2. Hu, B., Khara, P. & Christie, P. J. Structural bases for F plasmid conjugation and F pilus biogenesis in Escherichia coli. Proc. Natl. Acad. Sci. 116, 14222–14227 (2019).

3. Costa, T. R. D. et al. Structure of the Bacterial Sex F Pilus Reveals an Assembly of a Stoichiometric Protein-Phospholipid Complex. Cell 166, 1436–1444.e10 (2016).

4. Mazel, D. & Davies, J. Antibiotic resistance in microbes. C. Cell. Mol. Life Sci 56, 742–754 (1999).

5. Podschun, R. & Ullmann, U. Klebsiella spp. as nosocomial pathogens: Epidemiology, taxonomy, typing methods, and pathogenicity factors. Clinical Microbiology Reviews 11, 589–603 (1998).

6. Chen, L. et al. Carbapenemase-producing Klebsiella pneumoniae: molecular and genetic decoding. Trends Microbiol 22, 686–696 (2014).

7. Doumith, M. et al. Major role of pKpQIL-like plasmids in the early dissemination of KPC-type carbapenemases in the UK. J. Antimicrob. Chemother. 72, 2241–2248 (2017).

8. Chen, L. et al. Complete sequence of a blaKPC-2-harboring IncFIIK1 plasmid from a Klebsiella pneumoniae sequence type 258 strain. Antimicrob. Agents Chemother. 57, 1542–1545 (2013).

9. David, S. et al. Genomic analysis of carbapenemase-encoding plasmids from Klebsiella pneumoniae across Europe highlights three major patterns of dissemination. bioRxiv https://doi.org/10.1101/2019.12.19.873935 (2020). doi:10.1101/2019.12.19.873935

10. Tsai, Y.-K. et al. Klebsiella pneumoniae outer membrane porins OmpK35 and OmpK36 play roles in both antimicrobial resistance and virulence. Antimicrob. Agents Chemother. 55, 1485–93 (2011).

11. Fajardo-Lubián, A., Ben Zakour, N. L., Agyekum, A., Qi, Q. & Iredell, J. R. Host adaptation and convergent evolution increases antibiotic resistance without loss of virulence in a major human pathogen. PLoS Pathog. 15, e1007218 (2019).

12. Wong, J. L. C. et al. OmpK36-mediated Carbapenem resistance attenuates ST258 Klebsiella pneumoniae in vivo. Nat. Commun. 10, (2019).

13. Waksman, G. From conjugation to T4S systems in Gram-negative bacteria: a mechanistic biology perspective. EMBO Rep. 20, (2019).

14. Christie, P. J. The Mosaic Type IV Secretion Systems. EcoSal Plus 7, (2016).

15. Low, H. H. et al. Structure of a Type IV Secretion System. Nature 508, 550–553 (2014).

16. Cai, S. J. & Inouye, M. EnvZ-OmpR interaction and osmoregulation in Escherichia coli. J. Biol. Chem. 277, 24155–24161 (2002).

17. Poirel, L., Bonnin, R. A. & Nordmann, P. Genetic features of the widespread plasmid coding for the carbapenemase OXA-48. Antimicrob. Agents Chemother. 56, 559–62 (2012).

18. Potron, A., Poirel, L. & Nordmann, P. Derepressed transfer properties leading to the efficient spread of the plasmid encoding carbapenemase OXA-48. Antimicrob. Agents Chemother. 58, 467–71 (2014).

19. Finnegan, D. & Willetts, N. The Nature of the Transfer Inhibitor of Several F-like Plasmids. Mol. Gen. Genet. MGG 119, 57–66 (1972).

20. Maneewannakul, S., Kathir, P. & Ippen-Ihler, K. Characterization of the F plasmid mating aggregation gene traN and of a new F transfer region locus trbE. J. Mol. Biol. 225, 299–311 (1992).

21. Klimke, W. A. & Frost, L. S. Genetic analysis of the role of the transfer gene, traN, of the F and R100-1 plasmids in mating pair stabilization during conjugation. J. Bacteriol. 180, 4036–4043 (1998).

22. Klimke, W. A. et al. The mating pair stabilization protein, TraN, of the F plasmid is an outer-membrane protein with two regions that are important for its function in conjugation. Microbiology 151, 3527–3540 (2005).

23. Pérez-Mendoza, D. & De La Cruz, F. Escherichia coli genes affecting recipient ability in plasmid conjugation: Are there any? BMC Genomics 10, (2009).

24. Berger, C. N. et al. Citrobacter rodentium Subverts ATP Flux and Cholesterol Homeostasis in Intestinal Epithelial Cells In Vivo. Cell Metab. 26, 738–752 (2017).

