## Supplementary material for "OmpK36 and TraN facilitate conjugal transfer of the *Klebsiella pneumoniae* carbapenem resistance plasmid pKpQIL": Supplemtal data

### Supplementary Figures

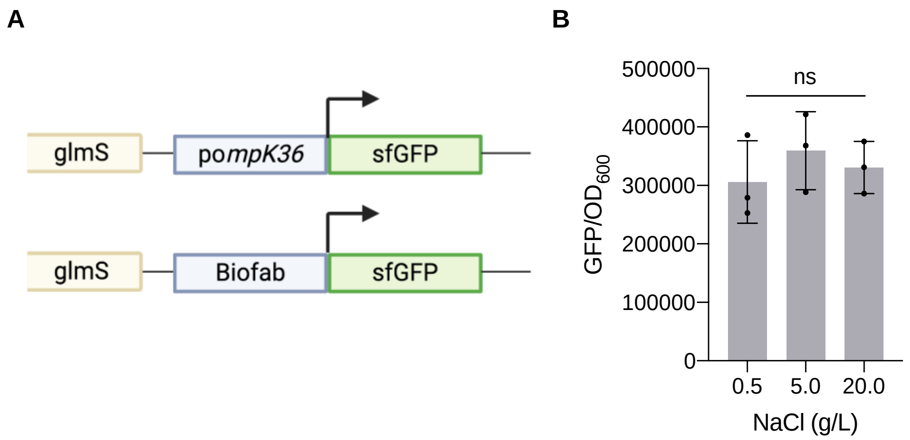


**Supplementary Figure 1. Assessing the effect of osmolality on OmpK36 expression.** (**A**) Schematic representation of the reporter system generated to measure promoter activity of OmpK36. A constitutive Biofab promoter-driven construct was generated to serve as a control for the system. Both constructs were inserted at the 3’ end of the *glmS* gene on the chromosome of ICC8001. (**B**) Activity of the Biofab promoter at different salt concentrations was measured by GFP emission. GFP emission was normalized to $OD_{600}$. ns = non-significant.

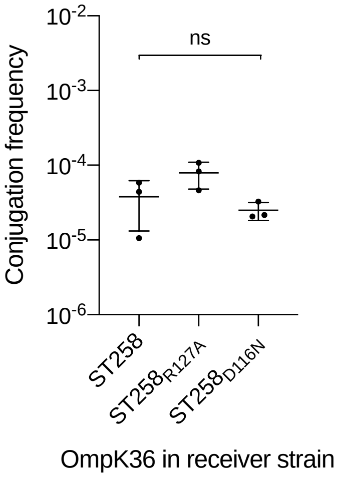


**Supplementary Figure 2. Assessing the effect of the salt bridge and charge on pKpQIL conjugation.** Conjugation frequency of pKpQIL was compared between recipients expressing $OmpK36_{ST258}$, $OmpK36_{ST258R127A}$ and $OmpK36_{ST258D116N}$. Conjugation frequency was determined as CFU transconjugants/ CFU recipients. ns = non-significant.

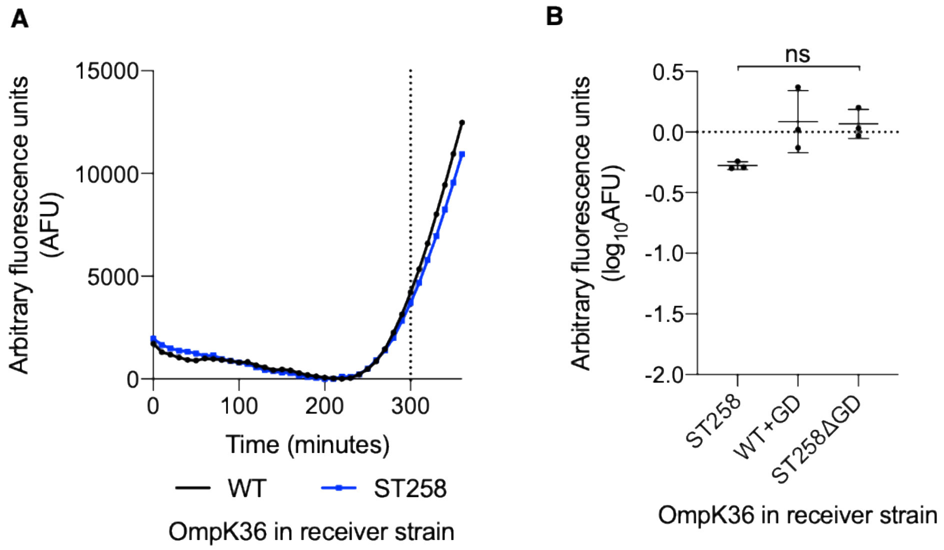


**Supplementary Figure 3. Real-time monitoring of pOXA-48a transfer.** (**A**) Plasmid transfer of an sfGFP-tagged pOXA-48a was monitored in real time by measuring GFP emission over 6 h into recipients expressing either OmpK36$_{WT}$ or OmpK36$_{ST258}$. Arbitrary fluorescence units (AFU) were normalised to the minimum GFP emission recorded for each conjugation mixture. (**B**) The log-fold difference in AFU recorded at *t* = 300 min respective to the OmpK36$_{WT}$-expressing recipient strain was calculated for recipients expressing various isoforms of OmpK36. ns = non-significant.

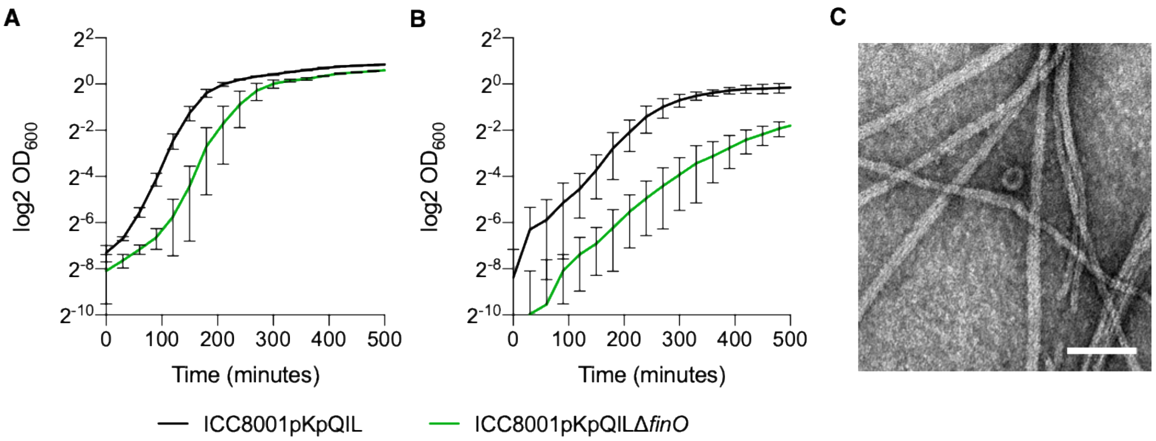


**Supplementary Figure 4. Characterisation of the derepressed donor strain.** (**A & B**) Growth of a donor strain, ICC8001, carrying pKpQIL or pKpQILΔ*finO* was compared in both rich (Luria Bertani) media (**A**) and minimal (M9) media (**B**) with glucose as the sole carbon source. (**E**) Negative stain of conjugative pili purified from ICC8001pKpQILΔ*finO*. Scale bar = 100 nM.

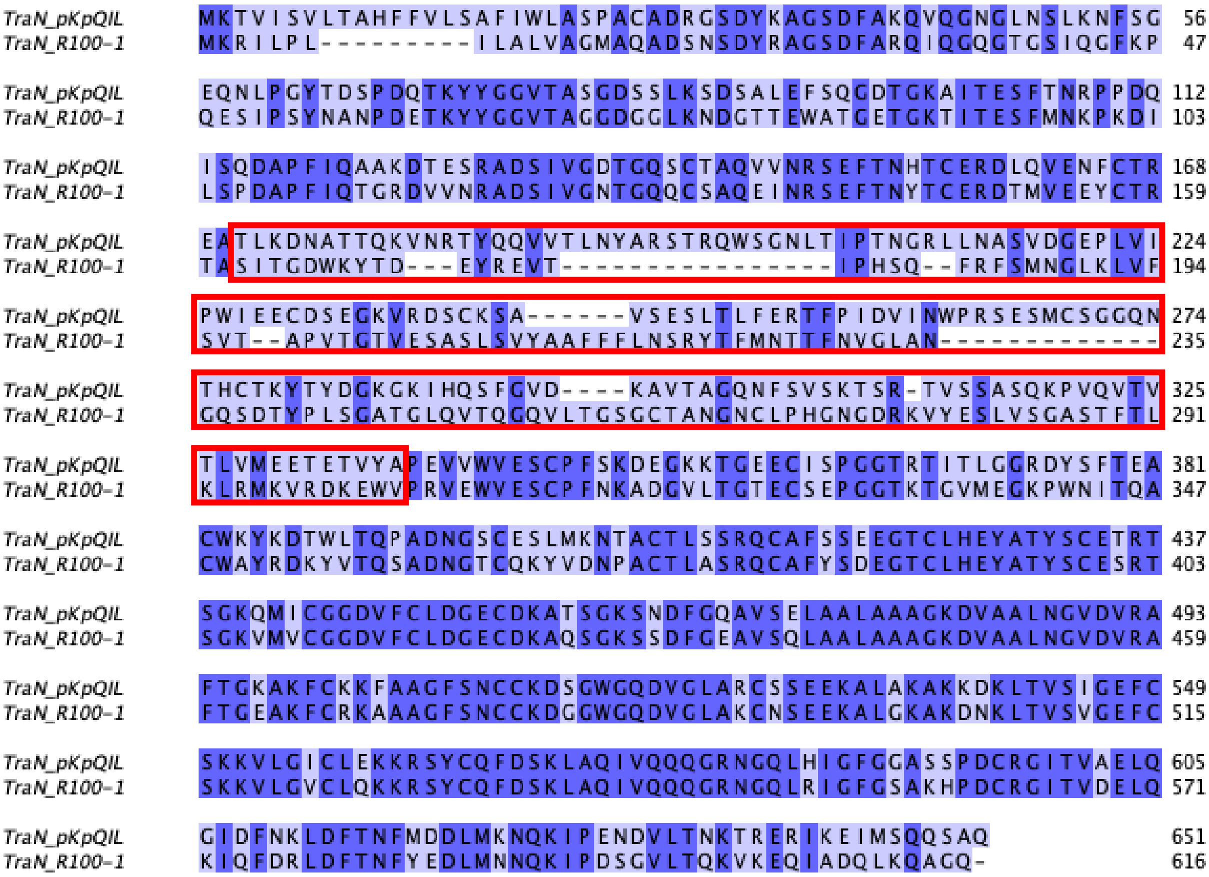


**Supplementary Figure 5. TraN sequence alignment.** The amino acid sequences of TraN from pKpQIL and R100-1 were aligned using Clustal Omega and visualized on Jalview. Amino acids highlighted in dark blue are conserved between both proteins and gaps are represented by dashes. The divergent region between aa 171 and aa 337 is outlined in red.

**Supplementary Table 1: Plasmid variants generated in this study**

| pICC no. | Plasmid |
|---|---|
| pICC4000 | pKpQIL-sfGFP |
| pICC4001 | pOXA-48a-sfGFP |
| pICC4002 | pKpQILΔ*finO* |
| pICC4003 | pKpQILΔ*finO*-sfGFP |
| pICC4004 | pKpQILΔ*traN*-sfGFP |
| pICC4005 | pKpQIL$_{\text{TraN_R100-1}}$-sfGFP |

**Supplementary Table 2: Strains used in this study.**

| No. | Strain | Description | Reference |
|---|---|---|---|
| S1 | ICC8001 | Conjugation donor strain. *K. pneumoniae* ATCC43816 serially passaged *in vitro* on Rifampicin (100 μg/ml) followed by two passages in BALB/c mice. | (Wong et al., 2019) |
| S2 | ICC8003 | Wild-type recipient strain. Isogenic derivative of ICC8001 expressing OmpK35$_{ST258}$ and OmpK36$_{WT}$ | |
| S3 | DH5α pKpQIL | *E. coli* transformant strain carrying pKpQIL | |
| S4 | DH5α pOXA-48a | *E. coli* transformant strain carrying pOXA-48a | |
| S5 | MDS42 R100-1 | *E. coli* strain carrying R100-1 | Gift from Fernando de la Cruz |
| S6 | CC118λpir | Maintains the R6K origin of pSEVA612S mutagenesis vectors | Lab collection |
| S7 | *E. coli* pRK2013 | Triparental mating helper strain | Lab collection |

**Supplementary Table 3: Vectors used in this study**

| No. | Vector | Resistance | Reference |
|---|---|---|---|
| V1 | pSEVA612S | Gentamicin | (Silva-Rocha et al., 2012) |
| V2 | pACBSR | Streptomycin | (Ruano-Gallego et al., 2015) |
| V3 | pSEVA-Kp-N2-GFP | Gentamicin | (Wong et al., 2019) |
| V4 | pSEVA-pOmpK36-GFP | Gentamicin | This study |
| V5 | pSEVA-OmpK36WT | Gentamicin | (Wong et al., 2019) |
| V6 | pSEVA-NHis-OmpK36 | Gentamicin | This study |
| V7 | pSEVA-OmpK36ST258 | Gentamicin | (Wong et al., 2019) |
| V8 | pET-28a (+) | Kanamycin | Lab collection |
| V9 | pSEVA-Kp-N2-lacI | Gentamicin | This study |
| V10 | pCSCMV-tdTomato | Ampicillin | Lab collection |
| V11 | pSEVA-Kp-N2-tdTomato | Gentamicin | This study |
| V12 | pSU2007::Tnlux | Kanamycin | Gift from Fernando de la Cruz (Pérez-Mendoza and De La Cruz, 2009) |
| V13 | pSEVA-pKpQILsfGFP | Gentamicin | This study |
| V14 | pSEVA-pOXA48sfGFP | Gentamicin | This study |
| V15 | pSEVA-pKpQILΔ*finO* | Gentamicin | This study |
| V16 | pSEVA-pKpQILΔ*traN* | Gentamicin | This study |
| V17 | pSEVA-pKpQIL$_{\text{TraN_R100-1}}$ | Gentamicin | This study |

**Supplementary Table 4: Primers used in this study**

| No. | Sequence 5' to 3' |
|---|---|
| P1 | GGTTAATAACATGCGTAAAGGCGAAGAG |
| P2 | GCATCCTATCTTGCAAAGGTCCGGTTG |
| P3 | TGCAAAGGTCCGGTTGAAATAGGGGTAAAC |
| P4 | GCATATAACAAACAGAGGGTTAATAACATGCGTAAA |
| P5 | CATCATCATCACCACCACGCTGAAATTTATAACAAAG |
| P6 | GCAGGCGCAGCAAATGCG |
| P7 | AACACCTACGGTTCTGACAACTTCCT |
| P8 | ATTCGGCGGCGACGGC |
| P9 | TTTAAGGAGGTAAAAAAAAAGTGGTGAATGTGAAACCAGTAAC |
| P10 | CTGGAAAGCGGGCAGTGATAAGGATCCAACAGGGTTCT |
| P11 | TAAGGATCCAACAGGGTTC |
| P12 | TTTTTTTTTACCTCCTTAAACTCC |
| P13 | TTTAAGGAGGTAAAAAAAAAATGGTGAGCAAGGGCGAG |
| P14 | AGAACCCTGTTGGATCCTTATTACTTGTACAGCTCGTCCATG |
| P15 | GAGGATACGTATGCGTAAAGGCGAAGAG |
| P16 | GGTATGGATGAACTGTACAAATAAGCAGATCAGT |
| P17 | GGCCTCGCGCGTCGAGAAAATTTATCAAAAAGAGTG |
| P18 | CAGGCTTGGAGGATACGTATGCGTAAAG |
| P19 | GGATTACCCTGTTATCCCTACAAACGCGAAGGCCGGTG |
| P20 | GATCGCTTGGCCTCGCGCGTCGAGAAAA |
| P21 | GTACAAATAAGCAGATCAGTTGGAAGAATTTG |
| P22 | GTTTGATAGTGGAATCTTGCATTACCCTGTTATCCCTATA |
| P23 | ATTACCCTGTTATCCCTATAC |
| P24 | GATTACCCTGTTATCCCTA |
| P25 | TGGCTGGGCGGTCGAGAAAATTTATCAAAAAGAG |
| P26 | GTATGGATGAACTGTACAAATAACGCTGATAAC |
| P27 | GGATTACCCTGTTATCCCTACGTACTCAACATCGGCGAAAGAG |
| P28 | TTTTCTCGACCGCCCAGCCACATCGTCC |
| P29 | GTACAAATAACGCTGATAACGTCTTTGCTG |
| P30 | CGTGAATGGCTGGATGCGATTACCCTGTTATCCCTATA |
| P31 | GGATTACCCTGTTATCCCTACCCGTGGTATTCCGGGAATATTC |
| P32 | GTAAATATAAAACAATTGCCTATCGTTCAGTTAATAAG |
| P33 | GGCAATTGTTTTATATTTACCCATTCCTGATAATTATACCTGGG |
| P34 | TATAGGGATAACAGGGTAATCGGCAACATCGTCTCCCC |
| P35 | GGATTACCCTGTTATCCCTACCGCCAGTTTATCGATAATCTG |
| P36 | GGATGAGGGAGGGCAGAAACCATGCTGC |
| P37 | GAGGGCAGAAACCATGCTGCCTAATAAAGAG |
| P38 | CTGAGCATGCCGCTATTCCATTACCCTGTTATCCCTATA |
| P39 | AGGACAGTAAACCATGCTGCCTAATAAAGAG |
| P40 | TACGTTTCATTTCTGCCCTCCCTCATCC |
| P41 | GAGGGCAGAAATGAAACGTATTTTACCTCTG |
| P42 | GCAGCATGGTTTACTGTCCTGCCTGTTTC |
